# Antibiotic inhibition of the *Plasmodium* apicoplast decreases haemoglobin degradation and antagonises dihydroartemisinin action

**DOI:** 10.1101/2021.10.31.466372

**Authors:** Emily M. Crisafulli, Amanda De Paoli, Madel V. Tutor, Ghizal Siddiqui, Darren J. Creek, Leann Tilley, Stuart A. Ralph

## Abstract

The World Health Organisation (WHO) recommends artemisinin (ART) combinations for treatment of uncomplicated *Plasmodium falciparum* malaria. Understanding the interaction between co-administered drugs within combination therapies is clinically important to prevent unintended consequences. The WHO guidelines recommend second line treatments that combine artesunate with tetracycline, doxycycline, or clindamycin—antibiotics that target the *Plasmodium* relict plastid, the apicoplast. In addition, antibiotics can be used simultaneously against other infectious diseases, leading to their inadvertent combination with ARTs. One consequence of apicoplast inhibition is a perturbation to haemoglobin uptake and trafficking—a pathway required for activation of ART derivatives. Here, we show that apicoplast-targeting antibiotics reduce the abundance of the catalyst of ART activation (free haem) in *P. falciparum*, likely through diminished haemoglobin digestion. We demonstrate antagonism between ART and these antibiotics, suggesting that apicoplast inhibitors reduce ART activation. These data have potential clinical implications due to the reliance on—and widespread use of—both ARTs and these antibiotics in malaria endemic regions.

## Introduction

Malaria remains one of the deadliest diseases affecting humankind, responsible for an estimated 409,000 deaths in 2019 [1]. Incidence and mortality rates remain high despite substantial efforts in drug and vaccine development, and the widespread use of bed nets and vector control programs in malaria endemic regions. Of particular concern in recent years has been the development of increasing resistance to the current frontline antimalarial, artemisinin (ART). The lack of comparably safe, effective, fast-acting, and affordable antimalarials in the drug pipeline signifies that efforts toward monitoring and managing parasite sensitivity to ARTs is of the upmost importance to prevent worsening of a global health emergency.

ART derivatives are recommended for use with a second partner drug as ART combination therapies, or ACTs, for the treatment of uncomplicated *Plasmodium falciparum* malaria [2]. The rationale for this is multifaceted, combining mitigation of risks against treatment failure—due to the very short in vivo half-life (∼1 h) of ART derivatives [3]—and against the development of drug resistance [4]—by hitting multiple drug targets within the parasite.

In some circumstances where preferred ART partner drugs are unavailable, the WHO recommends the use of artesunate (an ART derivative) plus either doxycycline or clindamycin [2]. There have also been calls to consider combining ART derivatives with antibiotics [5, 6], in part due to their dual anti-malarial and anti-bacterial activities. Currently, the WHO recommends treating patients presenting with suspected severe malaria with antibiotics in addition to malaria therapy—that is, until a bacterial infection can be excluded [2]. Use of these antibiotics in ACTs is appealing because many of them have anti-parasitic action through inhibition of the *Plasmodium* relict plastid, the apicoplast [7, 8]. Widespread use of these antibiotics in malaria endemic regions for malaria prophylaxis and the treatment of malaria or bacterial infections means that they may already be circulating in patients who seek treatment with ACTs for malaria. The deliberate as well as unintended use of these drugs in combination in the field makes it important to understand their modes of action, and possible interaction, within *P. falciparum* parasites.

The ability of these antibiotics to kill *Plasmodium* spp. relies on the bacterial origins of the apicoplast [8]. While inhibitors that directly target apicoplast metabolism generally cause immediate parasite death [9], others that target the organelle’s housekeeping functions—such as protein synthesis—cause a delayed death phenotype [10—12]. In delayed death, parasites survive the first cycle following treatment and it is not until the subsequent cycle that they cease proliferating and die. Isoprenoid synthesis is housed in the apicoplast, and antibiotic-treated parasites are depleted of the apicoplast-synthesised isoprenoid precursors required for the prenylation of trafficking machinery proteins, such as Rab proteins, resulting in a defect in the parasite’s uptake and digestion of haemoglobin [13, 14]. Haem released during haemoglobin digestion is predominantly responsible for the activation of ARTs via the cleavage of the endoperoxide bond [15, 16], so inhibition of this process may have implications for efficacy of ART treatment when these drugs are combined.

In this study, we aimed to assess the nature of the interaction between apicoplast-targeting antibiotics and ART derivatives. We hypothesised that these antibiotics would behave antagonistically with ARTs due to their inhibition of haemoglobin uptake and subsequent reduced release of free haem. We showed that delayed death antibiotics do indeed reduce the abundance of free haem available in *P. falciparum* for ART activation in the cycle after treatment. Although combinations of ART and antibiotics have previously been tested for interactions, those assays were designed to identify effects during the first cycle of treatment, which would miss effects that manifest at the time when delayed death antibiotics exert their biological impact. We instead tested for interactions in parasites that were treated with ART in the cycle after application of antibiotics. We found an antagonistic effect between these antibiotics and ART, presumably owing to the reduced activation of ART due to lower availability of free haem.

## Methods

### *Plasmodium falciparum* culture and synchronisation

*Plasmodium falciparum* 3D7 parasites were cultured as previously described [17]: at 2% (v/v) haematocrit in human O+ red blood cells (RBCs; Australian Red Cross Blood Service) and complete medium (RPMI-1640 with 25 mM sodium bicarbonate, 25 mM HEPES, 150 µM hypoxanthine, 20 µg/mL gentamicin (Sigma-Aldrich, G3632), and 0.5% (w/v) Albumax II (Thermo Fisher Scientific), pH 7.4); and maintained in malaria-mix gas (1% O_2_, 5% CO_2_, and 94% N_2_) at 37°C. Ring stage synchronisation (∼0−18 h p.i.) was achieved by a single treatment of infected RBCs with 5% (w/v) D-sorbitol (Sigma-Aldrich), unless otherwise indicated.

### 72 h in vitro single drug sensitivity assays

Synchronised ring stage 3D7 *P. falciparum* parasites (0.5% haematocrit; 1% parasitemia) were set up in V-bottom 96-well plates. Parasites were either: immediately treated with a dose gradient of atovaquone (Sigma-Aldrich, A7986), E-64 (Sigma-Aldrich, E3132), fosmidomycin (Sigma-Aldrich, F8682), proguanil (Sigma-Aldrich, G7048), or quinine (Sigma-Aldrich, 145904) in complete medium; or incubated for 24 h before treatment with various concentrations of dihydroartemisinin (DHA; Chem-Supply, D3793) or WR99210 (Jacobus Pharmaceutical) in complete medium. DHA-treated parasites were washed three times with complete medium 3 h post-drug treatment. Control wells containing parasites in the absence of drug or 100 µM chloroquine (Sigma-Aldrich, C6628) were prepared in parallel. All samples were prepared in triplicate.

72 h following set up, infected RBCs were incubated with lysis buffer (20 mM TRIS (pH 7.5), 5 mM EDTA, 0.008% (w/v) saponin, 0.08% (v/v) Triton X-100) and SYBR Green I (Thermo Fisher Scientific) for 1 h [18]. The fluorescent signal was detected using a FLUOstar Omega plate reader (BMG Labtech). Per cent survival was calculated by subtracting the background (chloroquine-treated) signal and normalising the data to the untreated control. GraphPad Prism (Version 9.1.2) was used to plot normalised data on XY scatter plots and calculate IC_50_s (presented as mean ± SEM).

### 120 h in vitro single drug sensitivity assays

Synchronised ring stage 3D7 *P. falciparum* parasites (0.5% haematocrit; 0.1% parasitemia) were set up in V-bottom 96-well plates. In the presence or absence of 5 µM geranylgeraniol (GGOH; Sigma-Aldrich, G3278), parasites were incubated in varying concentrations of clindamycin (Sigma-Aldrich, C5269) or doxycycline (Sigma-Aldrich, D3447) for 72 h, before the drugs were washed off with complete medium. Control wells were prepared (as described above, 72 h in vitro drug sensitivity assays) with chloroquine added to the positive control wells only at 72 h post-treatment. All samples were prepared in triplicate. At the 120 h time-point, infected RBCs underwent lysis, data acquisition, and analysis (as described above, 72 h in vitro drug sensitivity assays).

### Isobolograms

Synchronised ring stage 3D7 *P. falciparum* parasites (0.5% haematocrit; 72 h: 1% parasitemia, or 120 h: 0.1% parasitemia) were set up in V-bottom 96-well plates. Parasites were treated with dose gradients of two drugs, producing a 96-well plate where each well consisted of a unique combination (Fig. 1A). Various pairings of atovaquone, clindamycin, DHA, doxycycline, E-64, fosmidomycin, proguanil, quinine, and WR99210 were tested. The combination of delayed death drugs with DHA were done in both the presence and absence of 5 µM GGOH. As described above, for combinations that included DHA, parasites were washed three times with complete medium 3 h post-drug treatment, to mimic the short in vivo half-life of ARTs. Controls and treatment regimens for each combination were as previously described (72 h in vitro drug sensitivity assays; 120 h in vitro drug sensitivity assays; Fig. 2A to C, and S1A to C). All samples were prepared in duplicate.

**Fig 1.**
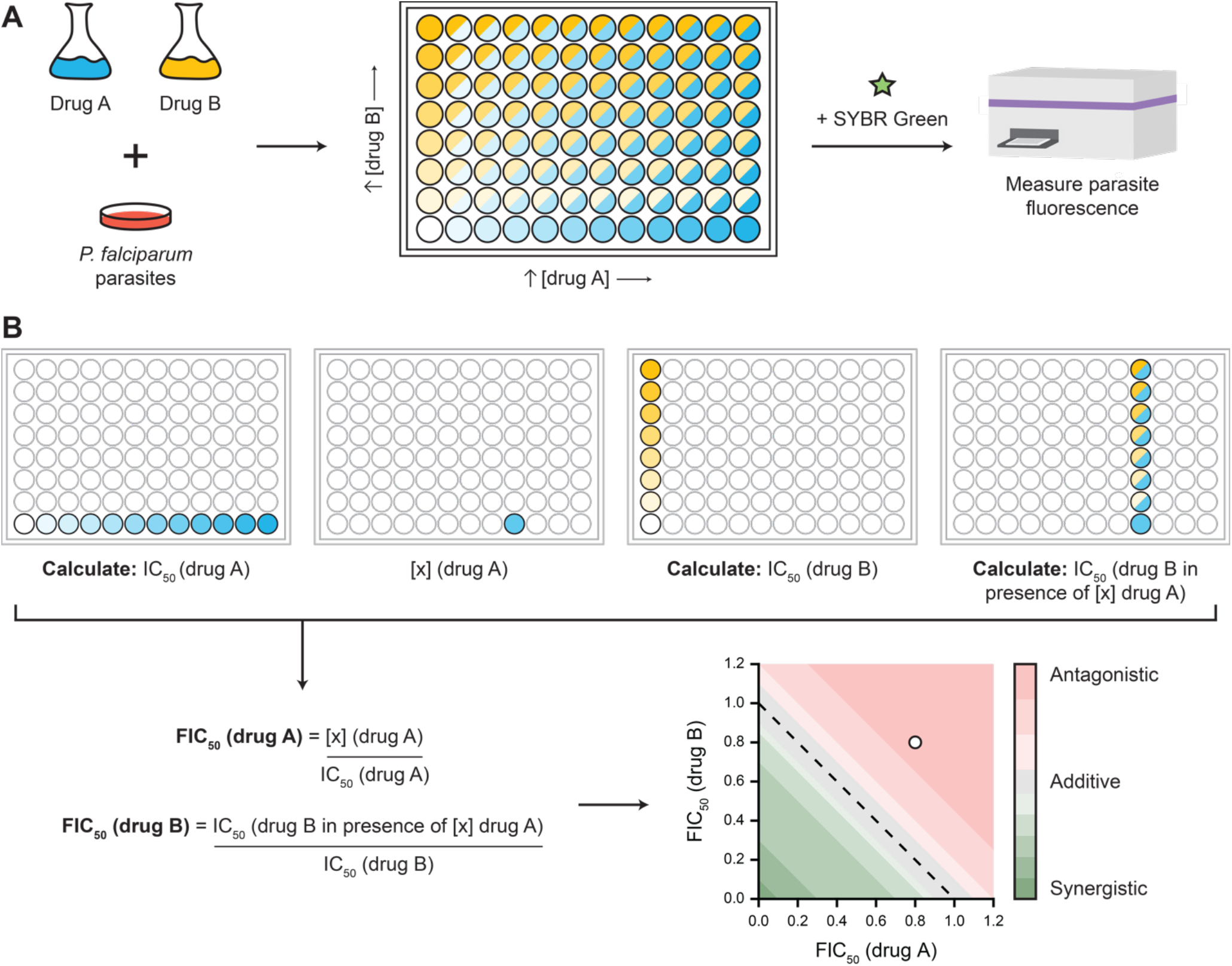
Drug interactions were determined by isobologram analysis. A) Schematic of the isobologram set-up pipeline, whereby 3D7 *Plasmodium falciparum* ring stage parasites were added to a V-bottom 96-well plate containing dose gradients of two drugs (blue, yellow). Following drug incubation, parasites were lysed and stained with SYBR Green for 1 h before fluorescent signal was detected using a microplate reader. B) Isobologram data analysis pipeline required calculation of fractional IC_50_s (FIC_50_s) to be plotted on an XY scatter plot. Drug combinations were evaluated to be synergistic (green), additive (grey), or antagonistic (red) based on previously defined thresholds [19].

**Fig 2.**
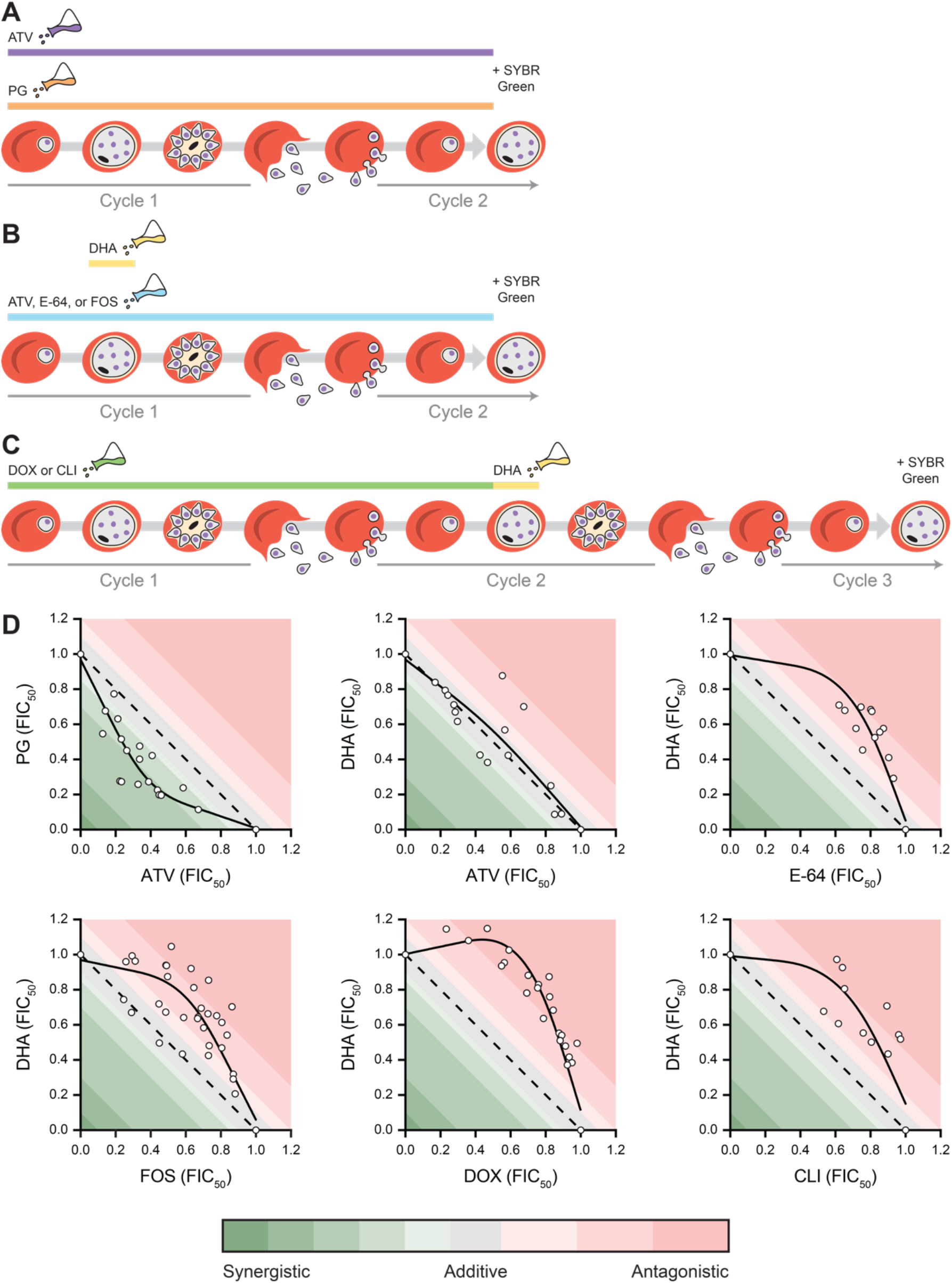
Normalised isobolograms demonstrating antagonism between apicoplast inhibitors and dihydroartemisinin (DHA). A—C) Schematic of the treatment regimen of 3D7 *Plasmodium falciparum* ring stage parasites in the set-up of D) isobolograms. Parasites were either: A) treated with dose gradients of atovaquone (ATV) and proguanil (PG) for 72 h; B) pre-treated for 24 h with ATV, E-64, or fosmidomycin (FOS); or C) pre-treated for 72 h with doxycycline (DOX) or clindamycin (CLI). Pre-treated parasites were pulsed with a dose gradient of DHA for 3 h. Parasites were lysed and stained with SYBR Green at A and B) 72 h or C) 120 h post-initial drug treatment. Fractional IC_50_s (FIC_50_s) are presented (n ≥ 3). Interaction thresholds as previously defined [19].

Parasites were lysed at the 72 or 120 h time-point (Fig. 2A to C, and S1A to C), data acquired, and per cent survival calculated for each data point (as described above, 72 h in vitro drug sensitivity assays). FIC_50_s were calculated [19] (Fig. 1B):

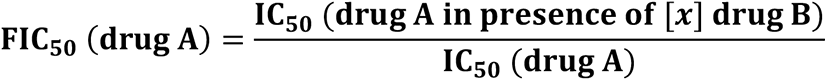

and GraphPad Prism (Version 9.1.2) used to plot data on XY scatter plots. ΣFIC_50_s were calculated to quantify the drug interaction (presented as mean ± SEM) [19]:

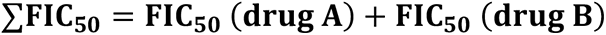

Interpretation of these values utilised previously defined thresholds [19]: < 0.1, very strong synergism; 0.1−0.3, strong synergism; 0.3−0.7, synergism; 0.7−0.85, moderate synergism; 0.85−0.9, slight synergism; 0.9−1.1, additive; 1.1−1.2, slight antagonism; 1.2−1.45, moderate antagonism; 1.45−3.3, antagonism; 3.3−10, strong antagonism; and > 10, very strong antagonism. GraphPad Prism (Version 9.1.2) was used to conduct unpaired *t*-tests with Welch’s corrections to test statistical significance of the ΣFIC_50_s.

### Haemoglobin fractionation

Ring stage 3D7 *P. falciparum* parasites were either: synchronised (as described above, *Plasmodium falciparum* culture and synchronisation) and treated with 3 µM fosmidomycin for 24 h; or subjected to two D-sorbitol treatments, 8 h apart, and parasites (∼8−18 h p.i.) subsequently treated with 1 µM doxycycline or 25 nM clindamycin for 68 h. Negative controls with the appropriate vehicle were set up in parallel. Following the incubation period, RBCs were lysed with 0.15% (w/v) saponin and parasite pellets washed three times with PBS and cOmplete™ mini EDTA-free protease inhibitor (Roche) at 4°C. Pellets were immediately frozen at −80°C until use.

The haemoglobin fractionation assay was adapted from [20, 21] with minor adjustments. Pellets were first resuspended in distilled water and sonicated for 5 min before addition of HEPES (pH 7.5; final concentration 0.1 M). Samples were centrifuged (6,200 x *g*, 20 min) and the pellets resuspended in distilled water before addition of 4% (w/v) sodium dodecyl sulfate (SDS; final concentration 2% (w/v)). The samples were sonicated for 5 min, then incubated at 95°C for 5 min to solubilise the free haem. A solution of HEPES, NaCl, and pyridine was added (final concentration 67 mM HEPES, 0.1 M NaCl, and 8.3% (v/v) pyridine) and samples centrifuged (6,200 x *g*, 20 min). The supernatant—containing the free haem fraction—was transferred to a clear, flat-bottom 96-well plate. The residual pellets were then resuspended in distilled water before solubilisation with NaOH (final concentration 0.15 M). They were sonicated for 15 min before a solution of HEPES, NaCl, and pyridine was added, as described above. These samples—corresponding to the haemozoin fraction—were transferred to the 96-well plate.

A blank (0.2 M HEPES, 0.3 M NaOH, 0.3 M NaCl, 0.3 M HCl, 4% (w/v) SDS, and 25% (v/v) pyridine) was included in triplicate for each study. The absorbance of each fraction was measured at 405 nm using an Ensight plate reader (PerkinElmer). Samples were blank adjusted and normalised to vehicle control. Data are presented as mean fold change compared to vehicle control ± SEM, and GraphPad Prism (Version 9.1.2) was used to perform one sample *t*-tests. All samples were prepared in triplicate.

## Results

### Apicoplast inhibitors behave antagonistically with the artemisinin derivative, dihydroartemisinin (DHA)

The Chou and Talalay isobologram method was used to test the efficacy of various drug combinations [22]. Specifically, we investigated whether apicoplast inhibitors and the ART derivative, dihydroartemisinin (DHA), behave antagonistically. Concentrations giving 50 per cent inhibition of growth of sorbitol-synchronised 3D7 ring stage *P. falciparum* (IC_50_ values) were first determined for individual drugs by means of a SYBR Green drug assay (Table S1). These were consistent with IC_50_ values from previous reports and allowed determination of dose gradients required to establish isobolograms [23–28]. For each isobologram, parasites were treated with varying combinations of two drugs across these gradients (Fig. 1A) at the relevant dosing regimens indicated (Fig. 2A to C, and S1A to C). The fraction of the IC_50_ concentration of each drug required to generate 50 per cent inhibition—the fractional IC_50_ (FIC_50_)—was then calculated for each dose held constant across the plate and plotted to form an isobologram (Fig. 1B). The shape of the isobologram and sum of the FIC_50_s (ΣFIC_50_s) were used to determine the type of drug interaction for each combination using previously defined thresholds [19].

To first validate this methodology, synchronised ring stage parasites were treated with the well-established synergistic combination of atovaquone and proguanil (Fig. 2A). We analysed growth at 72 h post-treatment (standard timing to assess first cycle death). The isobologram indicated a moderately synergistic interaction and produced a mean ΣFIC_50_ of 0.72 ± 0.04—concordant with previous reports [27, 29, 30] (Fig. 2D and 3).

Testing the interaction of delayed death drugs (which kill in the second cycle) and DHA (which kills in the first cycle) was a complex task, so we began by substituting the delayed death drug for an apicoplast inhibitor, fosmidomycin, that causes first cycle killing. Fosmidomycin directly blocks apicoplast metabolism by inhibiting 1-deoxy-D-xylulose-5-phosphate reductoisomerase, an enzyme involved in the isoprenoid biosynthetic pathway [9]. Like other apicoplast inhibitors, fosmidomycin perturbs haemoglobin uptake, albeit in the first cycle [13]. To specifically test the impact of the haemoglobin degradation defect, it was necessary to pre-treat synchronised ring stage 3D7 parasites with a dose gradient of fosmidomycin prior to DHA treatment. Further, to mimic the clinical scenario as closely as possible in an in vitro setting, we pulsed parasites with DHA for 3 h at the trophozoite stage, 24 h after pre-treatment with the apicoplast inhibitor (Fig. 2B). The experiment was designed to target the parasite life stage for which ART derivatives are most active [24, 31] and to mimic the short in vivo half-life of DHA [3]. Using this approach, fosmidomycin and DHA displayed moderate antagonism with a mean ΣFIC_50_ of 1.29 ± 0.05 (Fig. 2D and 3). This phenotype had been observed previously [32, 33], though using a different methodology that did not explore the clinically relevant half-life of DHA [3], nor fosmidomycin-induced haemoglobin trafficking defects [13]. The antagonism between fosmidomycin and DHA is concordant with the well-established antagonistic interaction between DHA and E-64 [31]—a cysteine protease inhibitor that prevents haemoglobin degradation, presumably reducing activation of DHA by free haem [34]. Consistent with this, we also saw a moderately antagonistic interaction between E-64 and DHA—with a mean ΣFIC_50_ of 1.36 ± 0.04 (Fig. 2D and 3)—using the approach described above (Fig. 2B).

As a control, we substituted the apicoplast inhibitor for atovaquone, an inhibitor of cytochrome *b* that was not predicted to interact with DHA. The mean ΣFIC_50_ of atovaquone and DHA was 1.00 ± 0.04, indicating a simple additive interaction and demonstrating that the unusual pre-treatment approach employed in our methodology did not contribute to the observed phenotype (Fig. 2D and 3).

A longer pre-treatment was necessary to test the interaction between delayed death drugs and DHA, as the haemoglobin trafficking defects observed when parasites are treated with these apicoplast inhibitors are evident only in the cycle subsequent to treatment [14]. Ring stage *P. falciparum* parasites were pre-treated with doxycycline or clindamycin for 72 h, to recreate the delayed death effect, and subsequently pulsed with DHA for 3 h at the trophozoite stage in the second cycle (Fig. 2C). Isobologram analysis of these data demonstrate an antagonistic and moderately antagonistic interaction between DHA and doxycycline (mean ΣFIC_50_ of 1.49 ± 0.02), and DHA and clindamycin (mean ΣFIC_50_ of 1.44 ± 0.10), respectively (Fig. 2D and 3). The ΣFIC_50_s of these combinations significantly increased—approximately 1.5-fold—from that of the additive interaction of atovaquone and DHA previously described (Fig. 3). This interaction with delayed death antibiotics was specific to DHA. The combination of delayed death drugs with WR99210—an inhibitor of dihydrofolate reductase—did not show the same antagonism (Fig. 3, and S1C and D). Mean ΣFIC_50_ values for doxycycline and clindamycin combined with WR99210 were 1.21 ± 0.05 and 1.16 ± 0.05, respectively (Fig. 3), suggesting that this cohort of apicoplast inhibitors have a selective effect on DHA activity. This interaction appeared to reverse when parasites were supplemented with exogenous geranylgeraniol (GGOH), a polyprenol that restores haemoglobin trafficking by permitting protein prenylation in the absence of isoprenoid biosynthesis [14] (Fig. S1B and D). However, the timing of these experiments was complex as GGOH-restored parasites survive longer but still die at a later stage that is yet to be thoroughly characterised [14], complicating attempts to quantify this interaction.

**Fig 3.**
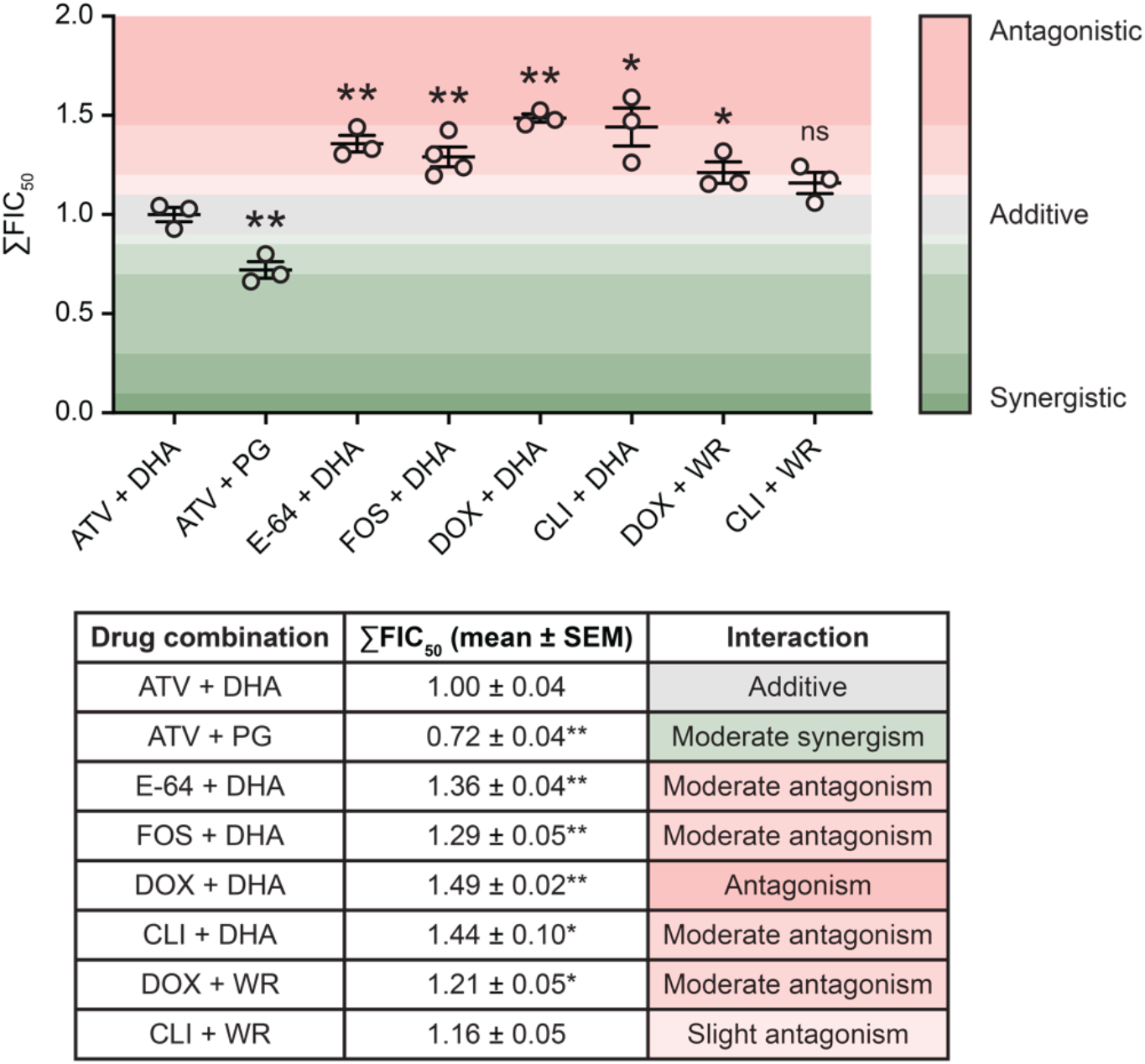
Mean ΣFIC_50_s demonstrating antagonism between apicoplast inhibitors and dihydroartemisinin (DHA). 3D7 *Plasmodium falciparum* ring stage parasites were either: treated with atovaquone (ATV) and proguanil (PG); pre-treated for 24 h with ATV, E-64, or fosmidomycin (FOS); or pre-treated for 72 h with doxycycline (DOX) or clindamycin (CLI). Pre-treated parasites were pulsed with a dose gradient of DHA for 3 h or WR99210 (WR) for 48 h. Parasites were lysed and stained with SYBR Green at 72 or 120 h. Data are presented as mean ± SEM (n ζ 3). Unpaired *t*-tests with Welch’s corrections were performed: *p ≤ 0.05,**p ≤ 0.01. Interaction thresholds as previously defined [19].

### Apicoplast inhibitors reduce the abundance of haemoglobin degradation by-products

To determine if the observed antagonistic interactions between apicoplast inhibitors and DHA were indeed due to reduced DHA activation, we sought to examine the effects of these antibiotics on the downstream products of haemoglobin digestion. Trafficking of haemoglobin is perturbed in parasites treated with fosmidomycin [13] or delayed death drugs [14], though the effect on by-products of haemoglobin digestion—free haem and haemozoin—has yet to be explored. We quantified the effects of fosmidomycin, doxycycline, and clindamycin on the abundance of these by-products in *P. falciparum* using a haem fractionation method, whereby pyridine is used to indirectly quantify haem products [20, 21]. The delayed death drugs, doxycycline and clindamycin, both significantly reduced haemozoin abundance in trophozoite parasites 72 h following treatment—a 1.3- (p = 0.0215) and 1.7-fold (p = 0.0416) reduction compared to the vehicle control, respectively (Fig. 4B). Fosmidomycin treatment caused a 1.5-fold reduction in haemozoin abundance in the same cycle as treatment. Though these latter changes were not statistically significant (p = 0.0854) (Fig. 4A), the magnitude of reduction is similar to that seen for other inhibitors of haemozoin formation assayed using the same methodology [20, 21]. While haemozoin appears reduced in abundance, the effect on free haem appears less stark for both fast-acting and delayed death apicoplast inhibitors (Fig. 4). However, this may be explained by it being a labile and toxic by-product of haemoglobin digestion that is quickly converted into chemically inert haemozoin by the parasite.

**Fig 4.**
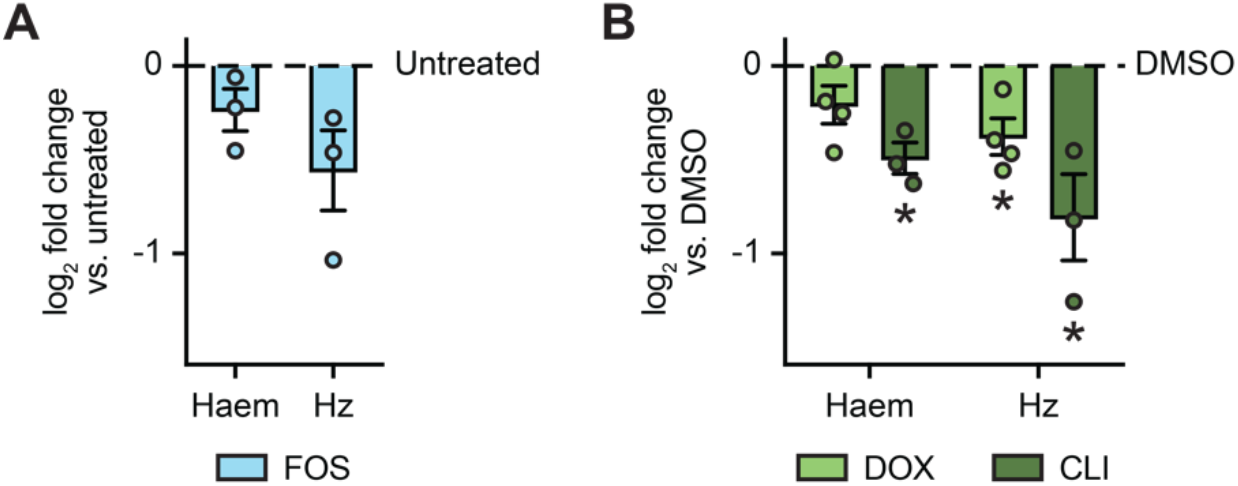
Apicoplast inhibitors reduce the abundance of haemoglobin degradation by-products. 3D7 *Plasmodium falciparum* ring stage parasites were treated with vehicle or: A) 3 µM fosmidomycin (FOS) for 24 h (∼24−42 h p.i.); or B) 1 µM doxycycline (DOX) or 25 nM clindamycin (CLI) for 68 h (∼76−86 h p.i.). Parasites were harvested and sequentially fractionated to determine the relative levels of free haem and haemozoin (Hz). Data were normalised to the vehicle control and are presented as mean log_2_ fold change ± SEM (n ≥ 3). One sample *t*-tests were performed: *p ≤ 0.05.

## Discussion

Combining drugs to enhance potency and reduce the risk of antimalarial resistance is a central plank in the strategies for malaria treatment. A notable example is the pairing of atovaquone with proguanil, a highly synergistic combination used for prophylaxis and treatment, frequently sold under the brand name Malarone [2, 29]. However, there are many documented examples of suboptimal antimalarial combinations [35], making choice of drug combinations key for effective treatment and reduced risk of drug resistance. Here, we demonstrate that antibiotics targeting the *P. falciparum* apicoplast behave antagonistically with the ART derivative, DHA, when the antibiotics are administered in the cycle before DHA. We propose that this interaction is underpinned by an antibiotic-mediated reduction in free haem that reduces the activation of DHA necessary for cytotoxicity (Fig. 5).

**Fig 5.**
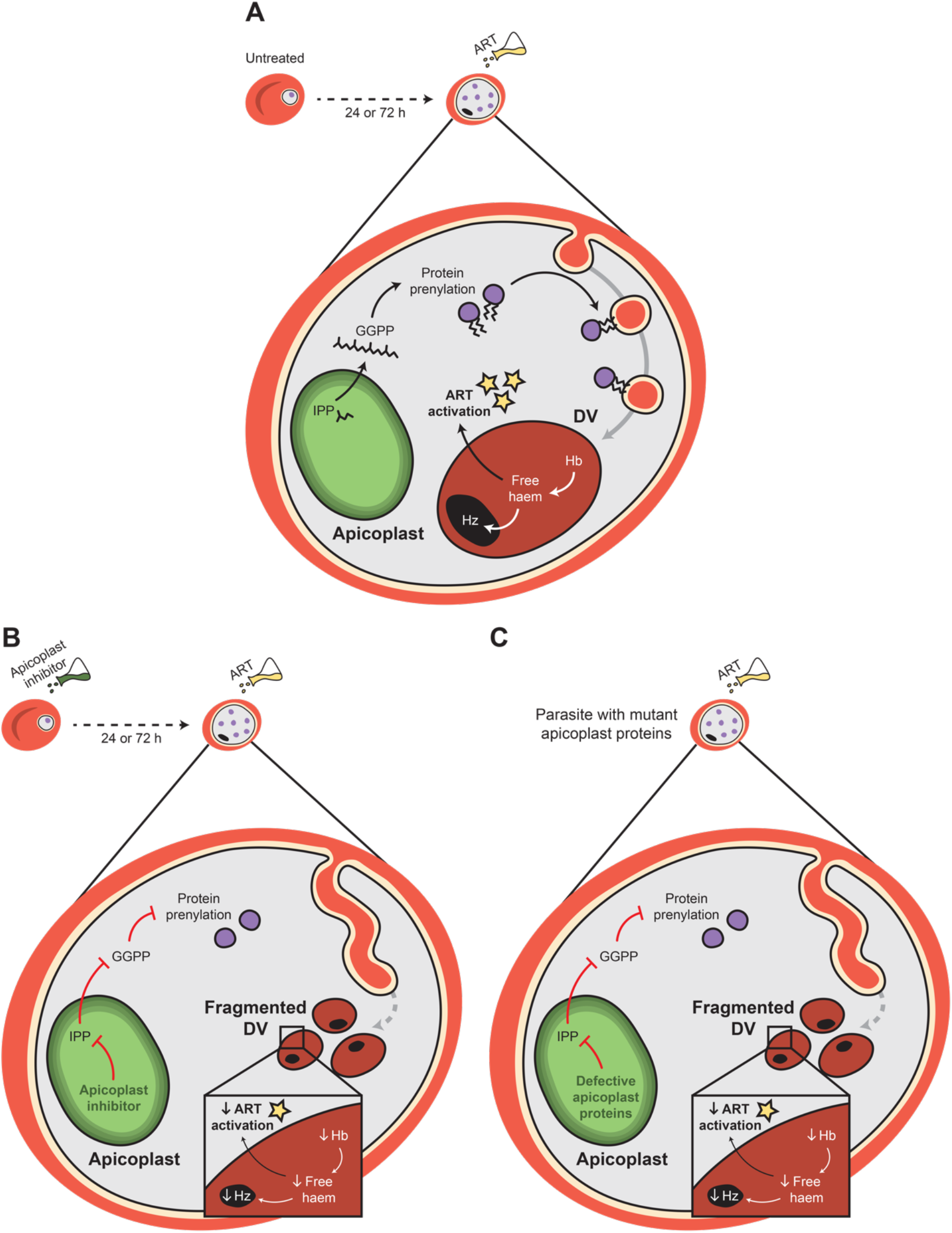
Model of apicoplast inhibition of isoprenoid biosynthesis decreasing haemoglobin (Hb) degradation and antagonising artemisinin (ART). A) In the absence of apicoplast inhibition, ART is activated by a product of Hb degradation—free haem—generated in the digestive vacuole (DV). Hb trafficking and degradation is dependent on the prenylation of trafficking proteins, a process that itself relies on production of the isoprenoid precursor, isopentenyl pyrophosphate (IPP), in the apicoplast. B) Apicoplast inhibition—both by fast-acting direct inhibitors of isoprenoid biosynthesis (24 h) and delayed death antibiotics (72 h)— reduces isoprenoid biosynthesis, preventing formation of geranylgeranyl pyrophosphate (GGPP), which forms the prenyl group on trafficking proteins required for Hb uptake and trafficking to the DV. In the absence of prenylated trafficking proteins, the cytostome becomes elongated and the DV fragmented. Hb degradation is reduced, reducing the abundance of free haem and haemozoin (Hz), and, subsequently, the activation of ART. C) Mutations in genes for apicoplast proteins reduce apicoplast metabolic processes, such as isoprenoid biosynthesis. In a similar fashion to drug inhibition, these mutations reduce prenylation and perturb haemoglobin uptake, such that there is less free haem available to activate ART.

Previous studies have mapped the effects of apicoplast inhibition in asexual *P. falciparum* parasites—from impeding IPP biosynthesis to downstream perturbation of haemoglobin uptake [13, 14, 36]. Isoprenoids have multiple cellular fates, though the proximate cause of parasite death through antibiotic treatment results from reduced prenylation of trafficking proteins, potentially through Rab proteins involved in haemoglobin trafficking to the DV [13, 14] (Fig. 5A). This prevents anchoring to vesicle membranes; and is associated with the aberrant uptake of haemoglobin from the host RBC and fragmentation of the DV [13, 14] (Fig. 5B). Further, delayed death drugs reduce the abundance of haemoglobin-like peptides in the cycle following treatment [14], consistent with our finding that free haem and haemozoin levels are lowered at this timepoint. Our data also demonstrate a reduction in abundance of these by-products by fosmidomycin—a fast-acting apicoplast inhibitor—suggesting that reduced haemoglobin degradation is an inevitable downstream effect of apicoplast inhibition in the asexual blood stages.

We hypothesise that this reduced release of free haem is the root cause of the antagonism between these antibiotics and DHA (Fig. 5). Consistent with this interpretation of the data is the antagonistic interaction observed between the cysteine protease inhibitor, E-64, and DHA—one that has been reported previously [31]. Indeed, a similar interaction has also been reported for another haemoglobin degradation inhibitor (the cysteine protease inhibitor, N-acetyl-L-leucyl-L-leucyl-L-norleucinal, or ALLN) [16]. In both studies, antagonism was attributed to a haem-mediated reduction in ART activation. This antagonistic interaction with apicoplast-inhibiting antibiotics may also extend to other ART-like compounds in the drug pipeline that similarly rely on free haem for activation—a notable example being ozonides [37, 38], the activation of which is directly inhibited by E-64 [39].

The antagonistic interaction between the fast-acting apicoplast inhibitor, fosmidomycin, and DHA described here is consistent with prior reports [32, 33]. By contrast, findings vary when testing the interaction between delayed death drugs and ART derivatives, although to our knowledge, such studies are restricted to analyses of interactions in the same cycle, which would ignore the impact of delayed death. For example, in vitro studies have reported interactions ranging from additivity [40—43] to synergy [42]; while there is a single in vivo report of synergy [44]. However, these studies were conducted using growth assays where the ART and antibiotic were administered at the same time and/or were terminated before the delayed death effect would have manifested. It is probable then that any effect captured results from the influence of secondary targets within the parasite, or only capture killing based on the very start of inhibition due to the antibiotic. Our experimental design specifically incorporates the delay in onset of the haemoglobin trafficking defect [14].

While combination treatment options would normally involve the simultaneous administration of ART with an antibiotic, several treatment regimens could mirror a scenario in which ART is present in the cell at a time where delayed death is relevant. One scenario that presents is the purposeful use of apicoplast inhibitors and ARTs in combination. The WHO currently recommends treating patients presenting with an unknown illness with antibiotics in addition to antimalarials—that is, until a bacterial infection can be excluded [2]. Antibiotics can also be used as partner drugs in ACTs: calls for these to be considered as frontline therapies have been made due to the added benefit that comes by concurrently treating any present bacterial infections prevalent in malaria endemic regions [5, 6]. While pre-treatment with antibiotics is not done purposefully in these instances, the need for multiple doses and the longevity of the antibiotic mode of action means that ultimately a similar “pre-treatment” scenario would eventuate, for example in the second or later dose of a combination therapy.

The widespread use of antibiotics in regions with high malaria transmission presents another— more complex—scenario whereby these drugs are combined inadvertently. Use of apicoplast-targeting antibiotics in mass drug administration efforts to treat bacterial infections in malaria endemic regions is one such situation—a notable example being administration of azithromycin for trachoma [45]. Azithromycin would consequently be circulating in patients seeking treatment for malaria infections with ACTs in these regions. There are also documented, and presumably many more undocumented, cases where non-compliance with a prescribed course of prophylactic doxycycline (which is rather commonplace [46—48]) results in a malaria infection [49]—an infection that will presumably be treated with an ACT. Both scenarios create conditions for possible suboptimal activation of the ART component of the ACT.

The many possible contributors (e.g. ART resistance) make it very difficult to deconvolve the root cause of any treatment failure and attribute it to this apicoplast drug interaction. However, clinical trial data—while presenting extremely varied reports of efficacy—suggest treatment failure is a very real possibility. A number of clinical trials report combinations of apicoplast inhibitors and ART derivatives that produced inferior cure rates and increased rates of recrudescence and treatment failure compared to other therapies [50–64] (Table 1).

**Table 1.**
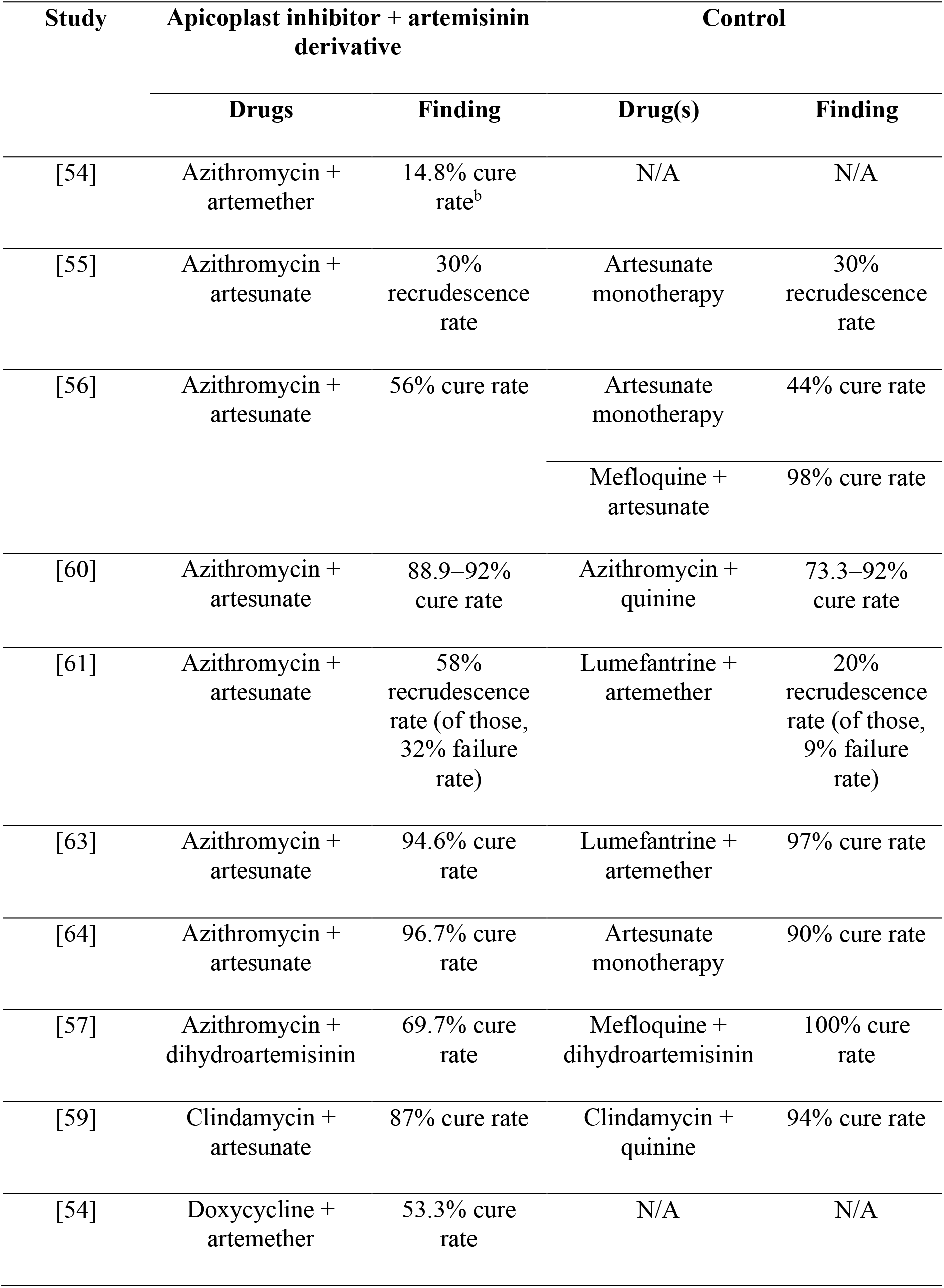
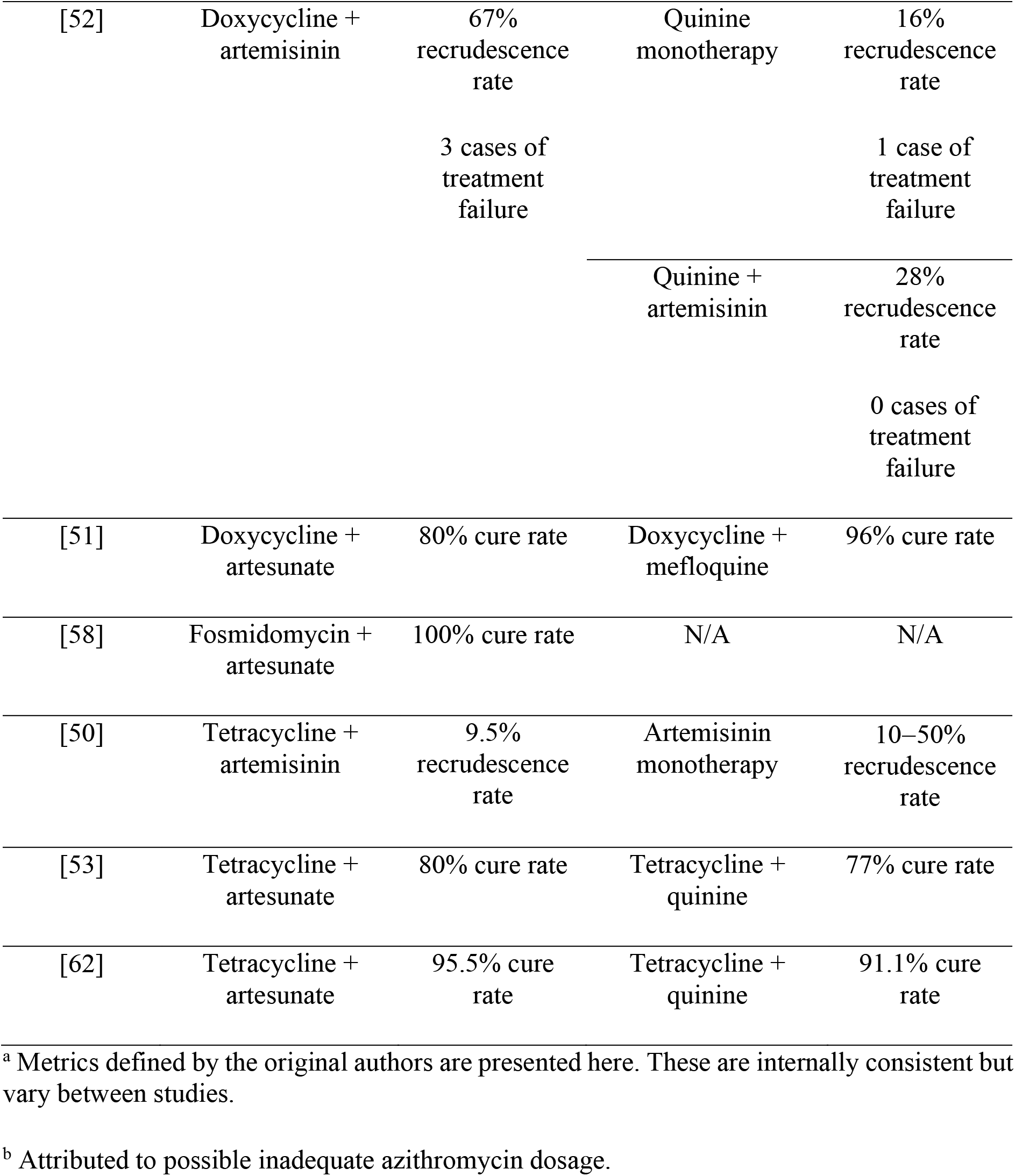
Summary of findings from clinical studies that combined apicoplast inhibitors with artemisinin derivatives^a^.

Arguably of greater global concern is that the combination of ART and apicoplast drugs could expose parasites to suboptimal concentrations of activated ART, conceivably worsening the already alarming spread of ART resistance and ART treatment failure that is currently occurring in South-East Asia [1]. There are several reports of modulation of ART sensitivity associated with mutations in or upstream of genes for apicoplast proteins, both in the field [65, 66] and in cultured parasites [67]. The significance of this is so far unclear, but one potential role could be through changes to apicoplast metabolism that impact isoprenoid synthesis and thus haemoglobin uptake (Fig. 5C). Given the almost ubiquitous use of ACTs throughout the malaria endemic world and the recent emergence of ART resistance in sub-Saharan Africa [68, 69], protecting against the rise of resistance elsewhere is key to avoid the worsening of malaria as a global health challenge.

The clinical uses of apicoplast-targeting antibiotics mean that they are often either purposefully or inadvertently used in combination with ART derivatives in the field. Although extrapolation of clinical relevance from in vitro data should be done carefully, these data flag potential concerns against combining ARTs with apicoplast-inhibiting antibiotics and reinforce the need to consider the molecular modes of action of any drugs used in combination in the field.

## Acknowledgments

We are grateful to Geoffrey I. McFadden, Christopher D. Goodman, Stanley C. Xie (University of Melbourne), Kit Kennedy (Weill Cornell Medicine), Matthew P. Challis (Monash Institute of Pharmaceutical Sciences), and Christina Spry (The Australian National University) for helpful discussions. We thank Jacobus Pharmaceutical for the kind gift of WR99210 and the Australian Red Cross Blood Service for donation of red blood cells for in vitro culturing of parasites.

This study was funded through grants from the Australian National Health and Medical Research Council (Grant #1181336, 1139884 and #1163235).

We also wish to acknowledge the Traditional Custodians of the lands on which this project was conducted, the Wurundjeri People of the Kulin nation.

