## Supplementary figure 1 for "Antibiotic inhibition of the *Plasmodium* apicoplast decreases haemoglobin degradation and antagonises dihydroartemisinin action"

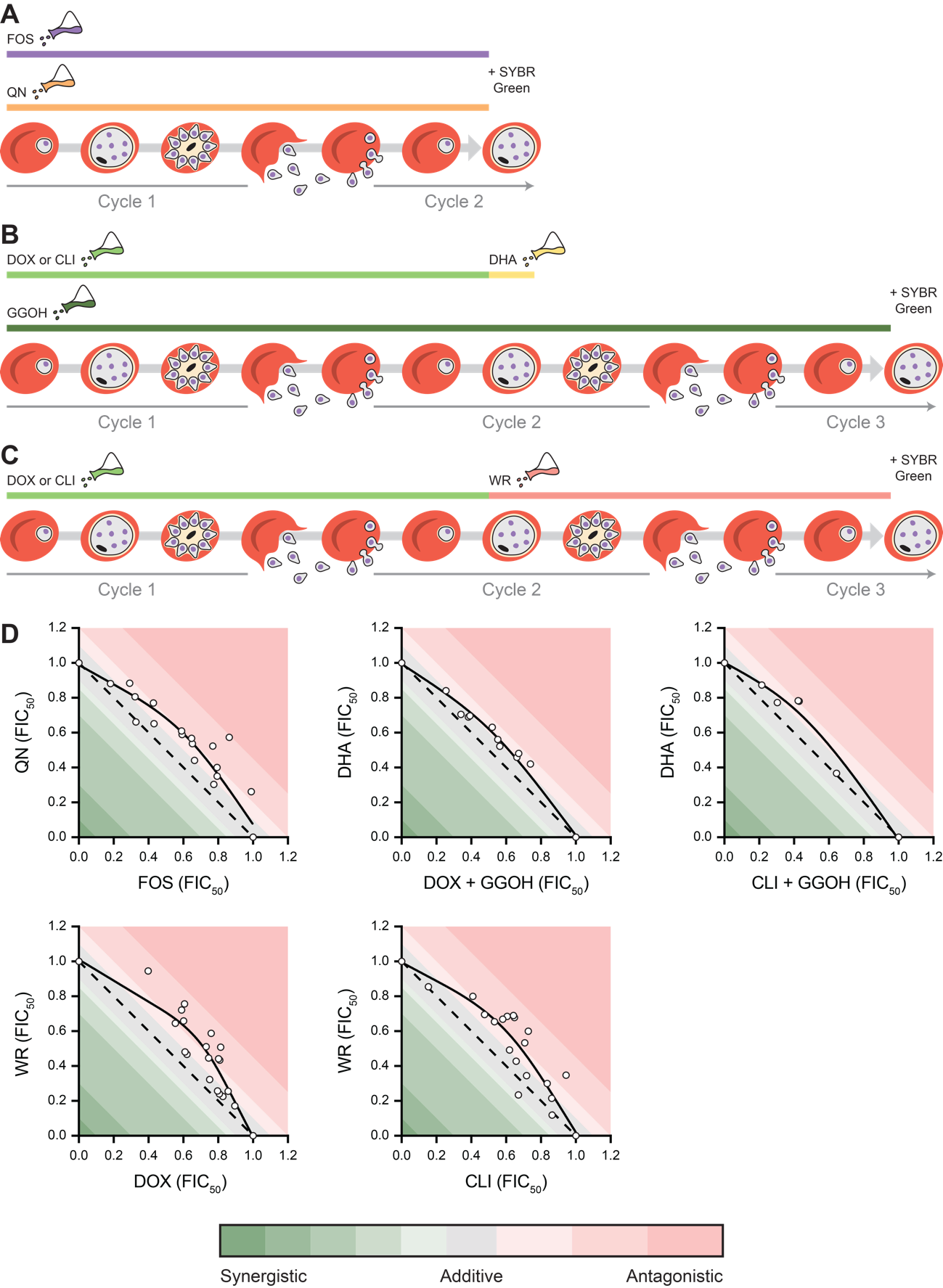

**Fig S1. Normalised isobolograms of various drug combinations.** A–C) Schematic of the treatment regimen of 3D7 *P. falciparum* ring stage parasites in the set-up of D) isobolograms. Parasites were either: A) treated with fosmidomycin (FOS) and quinine (QN); or B and C) pre-treated for 72 h with doxycycline (DOX) or clindamycin (CLI), in the B) presence or C) absence of 5 μM geranylgeraniol (GGOH). Pre-treated parasites were either: B) pulsed with a dose gradient of dihydroartemisinin (DHA) for 3 h; or C) treated with WR99210 (WR) for 24 h. Parasites were lysed and stained with SYBR Green at A) 72 h or B and C) 120 h. Data are presented as mean ± SEM (n ≥ 2). Interaction thresholds as previously defined [1].
