## Supplementary table 1 for "Antibiotic inhibition of the *Plasmodium* apicoplast decreases haemoglobin degradation and antagonises dihydroartemisinin action"

**Table S1. Inhibitory constants (IC_50_s) for *Plasmodium falciparum* parasites treated with various drugs.**

| **Drug** | **IC_50­_ (mean ± SEM)** | **Treatment regimen** | **Measurement time point post-treatment** |
| --- | --- | --- | --- |
| Atovaquone | 501.02 ± 60.85 pM | Rings; 72 h | 72 h |
| Clindamycin | 6.83 ± 1.67 nM | Rings; 72 h | 120 h |
| Clindamycin + GGOH^a^ | 21.35 ± 2.71 nM | Rings; 72 h (clindamycin); 120 h (GGOH) | 120 h |
| Dihydroartemisinin | 16.83 ± 1.51 nM | Trophozoites; 3 h | 48 h |
| Doxycycline | 747.07 ± 102.83 nM | Rings; 72 h | 120 h |
| Doxycycline + GGOH | 1.50 ± 0.30 μM | Rings; 72 h (doxycycline); 120 h (GGOH) | 120 h |
| E-64 | 6.27 ± 0.57 μM | Rings; 72 h | 72 h |
| Fosmidomycin | 1.10 ± 0.18 μM | Rings; 72 h | 72 h |
| Proguanil | 123.67 ± 18.39 nM | Rings; 72 h | 72 h |
| Quinine | 22.58 ± 1.38 nM | Rings; 72 h | 72 h |
| WR99210 | 623.41 ± 89.03 nM | Trophozoites; 48 h | 48 h |

^a^ Geranylgeraniol (GGOH).
